# Beer produced via hydrodynamic cavitation retains higher amounts of xanthohumol and other hops prenylflavonoids

**DOI:** 10.1101/176131

**Authors:** Rosaria Ciriminna, Lorenzo Albanese, Vita Di Stefano, Riccardo Delisi, Giuseppe Avellone, Francesco Meneguzzo, Mario Pagliaro

## Abstract

A fraction of the most valuable bioactive compounds in beer comes from hop’s polyphenols, mainly flavonoids, some of which are unique to inflorescences of that flowering plant. Although far from pharmacologically relevant concentrations, the intake of low doses of xanthohumol and related prenylflavonoids found in beer contributes to significant preventive actions with regards to certain diseases, such as cardiovascular and neurodegenerative, as well as to distinct chemopreventive effects on certain cancer types. Hence the efforts to explore both ingredients and methods able to enhance the concentration of such bioactive compounds in the final beers. A novel brewing method assisted by hydrodynamic cavitation was proved able, under suitable process conditions, to retain or generate higher amounts of xanthohumol, desmethylxanthohumol and 6-geranylnaringenin, extending recent evidence about different food and respective bioproducts, as well as adding to its distinct benefits to the brewing process.

## 1. Introduction

First detected in beer in 1999 by Stevens and co-workers in the US (Stevens, Taylor, & Deinzer, 1999), xanthohumol (XN, molecular formula C_21_H_22_O_5_) prenylated flavonoid secreted by hop (*Humulus lupulus*) inflorescences, is the most abundant polyphenol present in the hop lupulin glands and naturally found in the hard resins, as well as unique to hops, i.e. it has not yet been found in any other natural source (Almaguer, Schönberger, Gastl, Arendt, & Becker, 2014). Starting in 2008, XN has become the ingredient of numerous new dietary supplements (Stevens, 2012). Amidst several health benefits due to increasingly understood pharmacological profile (Liu et al., 2015), preclinical evidence proved XN’s action as antithrombotic (Xin et al., 2017), hepatoprotective (Weiskirchen, Mahli, Weiskirchen, & Hellerbrand, 2015), and anti-atherosclerotic (Hirata et al., 2017). While XN’s anti-carcinogenic properties, both alone and combined with other phenolic constituents extracted from malts and hops, have been revealed *in vitro* since some time (Ferk et al., 2010; Gerhäuser, 2005; Karabin, Hudcova, Jelinek, & Dostalek, 2015; Żołnierczyk et al., 2015), positive results of the first human intervention trials showed early clinical evidence (Pichler et al., 2017).

In general, pharmacologically relevant intakes of XN can only be achieved by oral administration of supplements, in doses from 16.9 mg/kg (action on glucose metabolism), up to 1 g/kg (anti-angiogenic action), still not showing any relevant toxicity, while beers usually contain no more than 0.2 mg/L (Liu et al., 2015). The amount of XN in alcoholic beers indeed varies, on average, between 0.10±0.07 mg/L in Ale-style beers and 0.3±0.03 mg/Lit in dark beers, depending on the hop varieties and the respective breeding conditions and age, as well as the brewing process (Almaguer et al., 2014; Rothwell et al., 2013; Stevens & Page, 2004). Indeed, xanthohumol and other prenylated hops’ flavonoids, such as desmethylxanthohumol (DMX, molecular formula C_20_H_20_O_5_), the latter polyphenol also unique to hops (Almaguer et al., 2014), undergo severe losses during the brewing process, due to incomplete extraction from hops in the wort, adsorption to insoluble malt proteins, and adsorption to yeast cells during fermentation. Furthermore, XN and DMX readily isomerize (cyclize) to the flavanones isoxanthohumol, and – due to two free *ortho* hydroxyl groups relative to C-1' in DMX – into a mixture of 6- and 8-prenylnaringenin, respectively, during the wort-boiling step of the brewing process (Karabin et al., 2015; Magalhães, Dostalek, Cruz, Guido, & Barros, 2008; Stevens, Taylor, Clawson, & Deinzer, 1999). As a result, contents of XN in final beers are usually 1% to 10% those of isoxanthohumol, with higher values in stout- and porter-style beers, while most beers contain no DMX due to thermal isomerization in the brew kettle (Stevens & Page, 2004). On the other hand, isoxanthohumol itself is endowed with anticarcinogenic properties, as well as, along with 8-prenylnaringenin, the latter being a conversion product of DMX in the brewing process and, after ingestion, a metabolic product of isoxanthohumol in the human liver and gut, with antioxidant activity, although at a lower efficiency than XN. As well, 8-prenylnaringenin has been identified as one of the most potent phytoestrogens identified so far (Karabin et al., 2015).

However, XN tops the other mentioned molecules as to biological and molecular activities (Venturelli et al., 2016), which explains the efforts to produce XN-enriched beers. Higher XN levels in beers can be achieved by means of late addition of hops to the boiling wort, addition of XN-enriched hops extracts, and the use of special malts, leading to concentrations as high as 3.5 mg/L (Magalhães et al., 2008), or even 10 mg/L without negatively affecting fermentation (Magalhães et al., 2011). Impacts of XN-enrichment on beers’ aroma, taste and shelf-life were found positive (Dresel, Vogt, Dunkel, & Hofmann, 2016).

Another interesting prenylated flavanone contained in hops is 6-geranylnaringenin (6-GN, molecular formula C_25_H_28_O_5_), also known as Bonannione A. It is found only in traces in hops and is mostly generated in the boiling wort from conversion of 3′-geranylchalconaringenin (3′-GCN, molecular formula C_25_H_28_O_5_), the latter being the only known geranylchalcone from hops (Stevens, Taylor, Clawson, et al., 1999). 6-GN displays significant antibacterial activity (Wang, Tan, Li, & Li, 2001), while little is known about its effects on cancer cells (Venturelli et al., 2016).

Although concentrations of XN and related prenylflavonoids in beers cannot achieve pharmacological levels, dietary intake of XN at the low doses commonly found in beers, under moderate consumption, has been suggested to possess distinct chemopreventive effects on certain cancer types, while much higher doses, despite lower XN’s bioavailability, are needed for cancer treatment (Blanquer-Rosselló, Oliver, Valle, & Roca, 2013). Indeed, recently it was proved that moderate beer consumption can be regarded as a healthy attitude due to preventive action with regards to certain diseases, such as cardiovascular and neurodegenerative, mainly as a result of the assumption of specific polyphenols from malts, grains and hops (de Gaetano et al., 2016), and to the overall antioxidant activity of some beers (Piazzon, Forte, & Nardini, 2010), which is higher than most wines and soft drinks (Queirós, Tafulo, & F. Sales, 2013). All the above, especially in craft beers that, lacking the clarification step, contrary to commercial products, retain most of naturally extracted polyphenols, in turn endowed with high bioavailability, thus leading to high antioxidant activity once ingested (Marques et al., 2017).

Among other substances, hop’s iso-α-acids were recently attributed specific and significant preventive action against the Alzheimer disease (Ano et al., 2017), as well as beer was estimated as one of the richest sources of silicon in the diet, which is important for the growth and development of bone and connective tissue (Casey & Bamforth, 2010).

The relevance of both the discovered healthy effects of moderate beer consumption, and the research on bioactive constituents of beers, arise from the fact that beer has become the worldwide most consumed alcoholic, with about 200 billion liters per year, being an integral part of the lifestyles of whole populations worldwide (Amienyo & Azapagic, 2016; Stack, Gartland, & Keane, 2016).

Recently, some of the authors introduced a new brewing process based on controlled hydrodynamic cavitation (HC) (Albanese, Ciriminna, Meneguzzo, & Pagliaro, 2017a). Operated directly on microbrewery scale (230 L), the new beer-brewing process affords a dramatic reduction of saccharification temperature, along with acceleration and increase of starch extraction, besides making traditional stages such as dry milling of malts and wort-boiling unnecessary at all. The latter comes along with significant time and energy savings, the latter estimated at a level well above 30%. In turn, avoiding wort-boiling was the combined result, achieved due to the cavitation process, of moderate-temperature sterilization, continuous removal of undesired volatile aromatic compounds such as dimethyl sulphide, and complete extraction and isomerization of hops’ α-acids before the onset of boiling. The complete utilization of hops’ α-acids in the HC-assisted beer-brewing process does not exhaust the matter of hopping, at least because, so far, the fate of XN and related prenylflavonoids was unknown. Investigating the resulting concentration of the latter important molecules in final beers produced by HC-assisted brewing is the subject of this study.

Shorter cleaning time, volumetric heating which prevents caramelization, overall simplification of both structural setup and operational management of brewing processes, no cavitational damage to the equipment arose, as well as no wort and beer oxidation, arose as further advantages.

Recently, an additional, important benefit brought by HC-assisted brewing was revealed by some of the authors: under certain cavitation regimes, the process allows achieving very low gluten content (<100 mg/L) or even gluten-free beers (<20 mg/L), starting from 100% barley malt as raw material (Albanese, Ciriminna, Meneguzzo, & Pagliaro, 2017b). Offering coeliac and gluten-intolerant people (the latter often unaware of the causes of their symptoms) with craft beers rid of toxic cereal proteins (collectively referred to as gluten), but fully retaining flavor and taste, can positively affect the public health. If important bioactive compounds, such as XN, can be retained, this would add to the healthy profile of the HC-assisted brewing process.

## 2. Materials and methods

### 2.1. Brewing unit

The beer-brewing unit powered by controlled HC, with total volume capacity around 230 L, performing the mashing and hopping stages, was comprehensively described in a previous study by some of the authors (Albanese et al., 2017a). Physics and chemistry of controlled HC in liquid media were introduced in other recent studies by some of the authors (Albanese et al., 2017b; Ciriminna, Albanese, Meneguzzo, & Pagliaro, 2017), as well as comprehensively analyzed, along with plenty of applications, by a growing number of recent studies (Carpenter et al., 2016; Šarc, Stepišnik-Perdih, Petkovšek, & Dular, 2017).

Fig. 1 shows the experimental installation, described in detail in section 2.1 of the previous study by the authors (Albanese et al., 2017a), along with the calculation, meaning and limitations of the basic HC metric, *i.e.* the Cavitation Number (CN). Here, it suffices recalling the reference to the description of the cavitation reactor in the form of a Venturi tube (Albanese, Ciriminna, Meneguzzo, & Pagliaro, 2015). All the tests ran in brew-in-the-bag (BIAB) mode, using the malts caging vessel shown in Fig. 1; however, hops were allowed circulating in any process.

**Figure 1.**
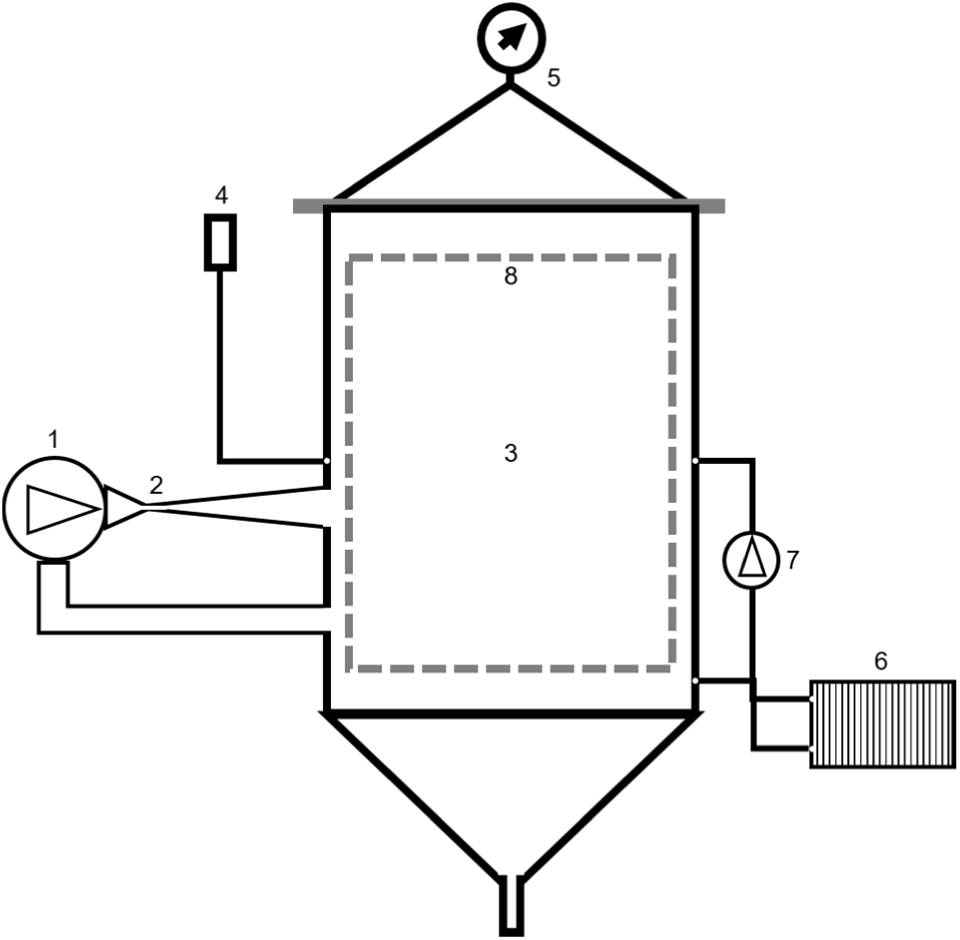
Simplified scheme of the experimental HC-based installation. 1 – centrifugal pump, 2 – HC reactor, 3 – main vessel, 4 – pressure release valve, 5 – cover and pressure gauge, 6 – heat exchanger, 7 – circulation pump, 8 – malts caging vessel. Other components are commonly used in state-of-the-art hydraulic constructions.

For comparison purposes, a parallel brewing test was performed by means of a conventional Braumeister (Ofterdingen, Germany) model 50 L brewer, equipped with a cooling serpentine (model Speidel stainless steel wort chiller for 50-liter Braumeister) and fully automatic brewing control (temperature, time and recirculation pumps).

### 2.2. Analytical instruments and methods

Along with thermometer and manometer sensors onboard the main production unit, few specialized off-line instruments were used to measure the chemical and physiological properties of wort and beer, described in full detail in a previous study by some of the authors (Albanese et al., 2017a). Few of those instruments are described herein below.

Physico-chemical and physiological parameters of finished beers were measured by means of a 6-channel photometric device (CDR, Firenze, Italy, model BeerLab Touch). In particular, fermentable sugars (0.1 to 150 g/L of maltose, resolution 0.01 g/L), alcohol content (0-10% in volume, resolution 0.1%), bitterness on the International Bitterness Unit (IBU) scale (5 to 100, resolution 0.1) (Lajçi, Dodbiba, & Lajçi, 2013), color on the European Brewery Convention (EBC) scale (1 to 100, resolution 1) and on the Standard Reference Method (SRM) scale (0.5 to 50, resolution 0.1). All reagents were of analytical grade.

The power and electricity consumed by the three-phase centrifugal pump used in the HC device shown in Fig. 1 were measured by means of a three-phase digital power meter (IME, Milan, Italy, model D4-Pd, power resolution 1 W, energy resolution 10 Wh, accuracy according to the norm EN/IEC 62053-21, class 1).

The instruments and methods specific to this study are described below.

Liquid chromatography coupled to mass spectrometry is now a well-established methodology for food analysis (Di Stefano et al., 2012). Hence, we have used LC-MS to assess the amount of xanthohumol, and of two other prenylated flavanones, desmethylxanthohumol, and 6-geranylnaringenin, in four different beer samples.

The structural formulas of the above-mentioned prenylflavonoids investigated in beer samples are depicted in Fig. 2.

**Figure 2.**
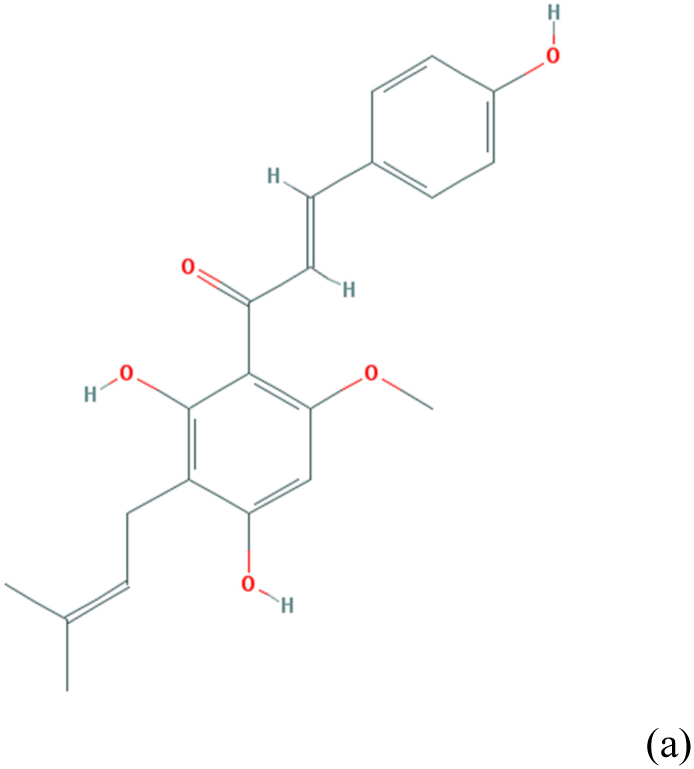
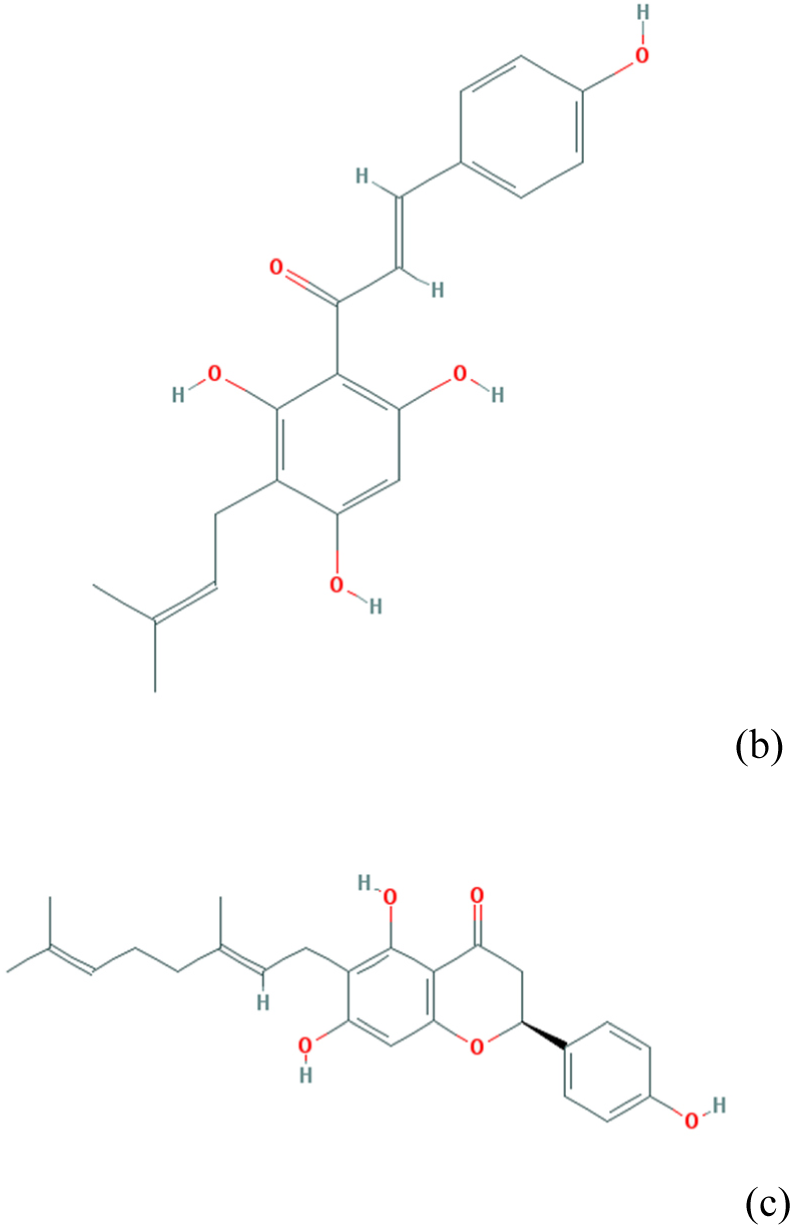
Structural formulas of the prenylflavonoids investigated in beer samples obtained via hydrocavitation: xanthohumol (a), desmethylxanthohumol (b) and 6-geranylnaringenin (c). Pictures drawn from PubChem (https://www.ncbi.nlm.nih.gov/pccompound).

The beer samples were filtered over PTFE syringe filter 0.45 μm. Xanthohumol (3’-[3,3-dimethyl allyl]-2’,4’,4-trihydroxy-6’-methoxychalcone) and related prenylflavonoids, *i.e.* desmethylxanthohumol and 6-geranylnaringenin, were identified by ultra-high performance liquid chromatography, heated electrospray, and mass spectrometry (UHPLC-HESI-MS). UHPLC analysis was performed using a Dionex Ultimate 3000 System (Dionex Softron GmbH, Germering, Germany) equipped with an autosampler controlled by Chromeleon 7.2 Software (Thermo Fisher Scientific, Bremen, Germany) coupled to a photodiode array detector (Thermo Fisher). A UHPLC column (Phenomenex Luna C18(2) 50x1mm, 2.5μ) was used for separation of the selected compounds at 30°C. The mobile phases used were 0.1% formic acid in water (A) and ACN with 0.1% formic acid (B). The gradient elution program was: 0-3 min 5% B; 3-30 min linear increase to 95% B, 30-35 min 95% and 35-40 min B returning to the initial conditions until full stabilization. The injection volume was 1 μl and the flow rate 50 μl min^-1^. MS detection was performed using a Q-Exactive accurate-mass spectrometer (Thermo Scientific, Bremen, Germany). The HESI parameters were set using negative ion mode with spectra acquired over a mass range from *m/z* 100-800.The optimum values of HESI-MS parameters were: sheath gas flow at 25 arbitrary units; auxiliary gas unit flow at 6 arbitrary units; capillary temperature at 320 °C; auxiliary gas heater temperature at 150 °C; spray voltage at 3.0 kV; and S lens radio frequency level set at 50. The automatic gain control was set with a maximum injection time of 200 ms. HESI-MS spectra yield single deprotonated ion, [M-H]^-^, at the same time as the mode FULL-SCAN and t-SIM (targeted Selected Ion Monitoring), to increase sensitivity. The total UHPLC-HESI-MS method run time was 40 min. Detection of xanthohumol was based on calculated exact mass and on retention time of standard compound. Desmethylxanthohumol and 6-geranylnaringenin were assessed only on the basis of calculated exact mass with data evaluated by Quan/Qual browser Xcalibur 3.0 (Thermo Fisher Scientific, San Jose, CA, USA). The linearity of the MS response was verified with solutions containing xanthohumol standard at seven different concentration levels over the range from 0.005 to 1 ppm.

Xanthohumol (purity ≥ 99%) was purchased by Extrasynthese, France. A stock standard solution was prepared at a concentration of approximately 0.1 mg mL^-1^ in 80:20 MeOH/H_2_O (v/v). The other standard solutions (at 1.0, 0.5, 0.2, 0.05, 0.02, 0.01, 0.005 ppm) were prepared by dilution of the stock solution. Calibration curves were constructed by injecting each standard solution at each concentration level in quadruplicate. The peak areas were calculated and plotted against the corresponding concentrations of the standard compound using linear regression (least squares) to generate the standard curve. Content of desmethylxanthohumol and 6-geranylnaringenin in samples were evaluated using xanthohumol as a reference compound.

### 2.3. Brewing ingredients

Pilsner or Pale were used as the base barley malts in all the performed tests, along with smaller fractions of Cara Pils and Cara Hell. Among the hops, different combinations of pelletized German Perle (hereinafter, Perle), Saaz and German Hersbrucker (hereinafter, Hers) were used. Table 1 shows the average concentration of the three prenylflavonoids under study in the used hops, drawn from available sources (Dresel et al., 2016; Stevens, Taylor, & Deinzer, 1999).

**Table 1.** Average concentration of xanthohumol, desmethylxanthohumol and 6-geranylnaringenin in the used hops.

| **Hop** | **Xanthohumol<sup>a)</sup>**<br>(mg/100g) | **Desmethyloxanthohumol<sup>a)</sup>**<br>(mg/100g) | **6-geranylnaringenin<sup>b)</sup>**<br>(mg/100g) |
| --- | --- | --- | --- |
| Perle | 1150 | 4.046 | 0.018 |
| Hers<sup>c)</sup> | 386.6 | 2.522 | 0.018 |
| Saaz | 1053 | 0.674 | 0.018 |

a) Source: (Dresel et al., 2016).
b) Source: (Stevens, Taylor, & Deinzer, 1999). Concentration of 6-GN is extremely low in hops and differences between hop's types are unknown, it being generated in the boiling wort from conversion of hops' geranylchalcone 3'-geranylchalconaringenin.
c) Hers = Hallertau Hersbrucker hop.

Fermentation was activated by means of the dry yeast strain Safale US-05, requiring temperature between 15°C and 24°C and maximum alcohol content 9.2%, used in any test in the identical proportion of 67 g per 100 L.

### 2.4. Brewing tests

The four different beer samples were prepared as described in a previous study by some of the authors (Albanese et al., 2017a), to which reference is made for further insight. Different absolute and relative amounts of hops were used. Table 2 summarizes few basic features of the brewing tests that, as stated in Section 2.1, ran in BIAB mode, yet with cavitating hops. In test C10, contrary to the others, the wort fermented into the same processing device shown in Figure 1, instead of traditional, separate fermenters. One of the tests (B1) was performed by means of the traditional Braumeister model 50 L brewer, described in Section 2.1.

**Table 2.** Beer production tests, ingredients, conditions, and basic features of final beers.

| Test ID | Production unit<sup>a)</sup> | Net volume (L) | Malts | Hops<sup>b)</sup> | Fermentation<sup>c)</sup> | Beer color<sup>d)</sup> (EBC / SRM) | Alcohol<sup>d)</sup> (%vol) | IBU<sup>d)</sup> |
| --- | --- | --- | --- | --- | --- | --- | --- | --- |
| B1 | B-50 | 50 | Pilsner 6.25 kg<br>Cara Pils 0.9 kg<br>Cara Hell 0.65 kg | Perle 0.025 kg<br>Hers 0.075 kg<br>Saaz 0.05 kg | Standard | 6 / 3.2 | 3.2 | 21.7 |
| C6 | HC | 170 | Pilsner 25 kg<br>Cara Pils 3.6 kg<br>Cara Hell 2.6 kg | Perle 0.3 kg<br>Hers 0.2 kg<br>Saaz 0.4 kg | Standard | 6 / 2.8 | 5.0 | 38.9 |
| C7 | HC | 170 | Pilsner 28.5 kg<br>Cara Pils 2.5 kg | Hers 0.6 kg<br>Saaz 0.5 kg | Standard | 5 / 2.7 | 3.4 | 30.9 |
| C10 | HC | 170 | Pale 26 kg<br>Cara Pils 3 kg | Perle 0.2 kg<br>Hers 0.1 kg | Installation | 5 / 2.5 | 3.9 | 20.0 |

a) B-50 = Braumeister model 50-liters; HC = experimental installation shown in Figure 1 – variant with malts caged in a cylindrical vessel (BIAB).
b) Hops cavitating in any test.
c) Standard = fermentation in cylinder-conical vessel 200 L. Installation = fermentation performed into the experimental installation HC.
d) Beer color, alcohol content and IBU were measured 15±2 days after bottling.

Color and alcohol levels shown in Table 2 match the Blonde Ale beer style for all samples, as expected from the respective recipes (Priest & Stewart, 2006).

Fig. 3(a-d) summarize the brewing processes in terms of temperature and cavitation number against consumed energy, for HC-assisted tests C6, C7 and C10, and in terms of temperature against time for conventional test B1. Data points start at the time of mashing-in (insertion of malt), while mashing-out and hops input are highlighted in the charts, as well as yeast pitching in test C10, when HC was activated also during the early fermentation stage (*i.e.*, after yeast pitching), at temperatures around 27°C and during about 4 hours. Fermentation lasted 12 days in test C6, 7 days in test C7, and 8 days in test C10, as also shown in Fig. 3 of a previous study by some of the authors (Albanese et al., 2017b), while it lasted 7 days in test B1.

**Figure 3.**
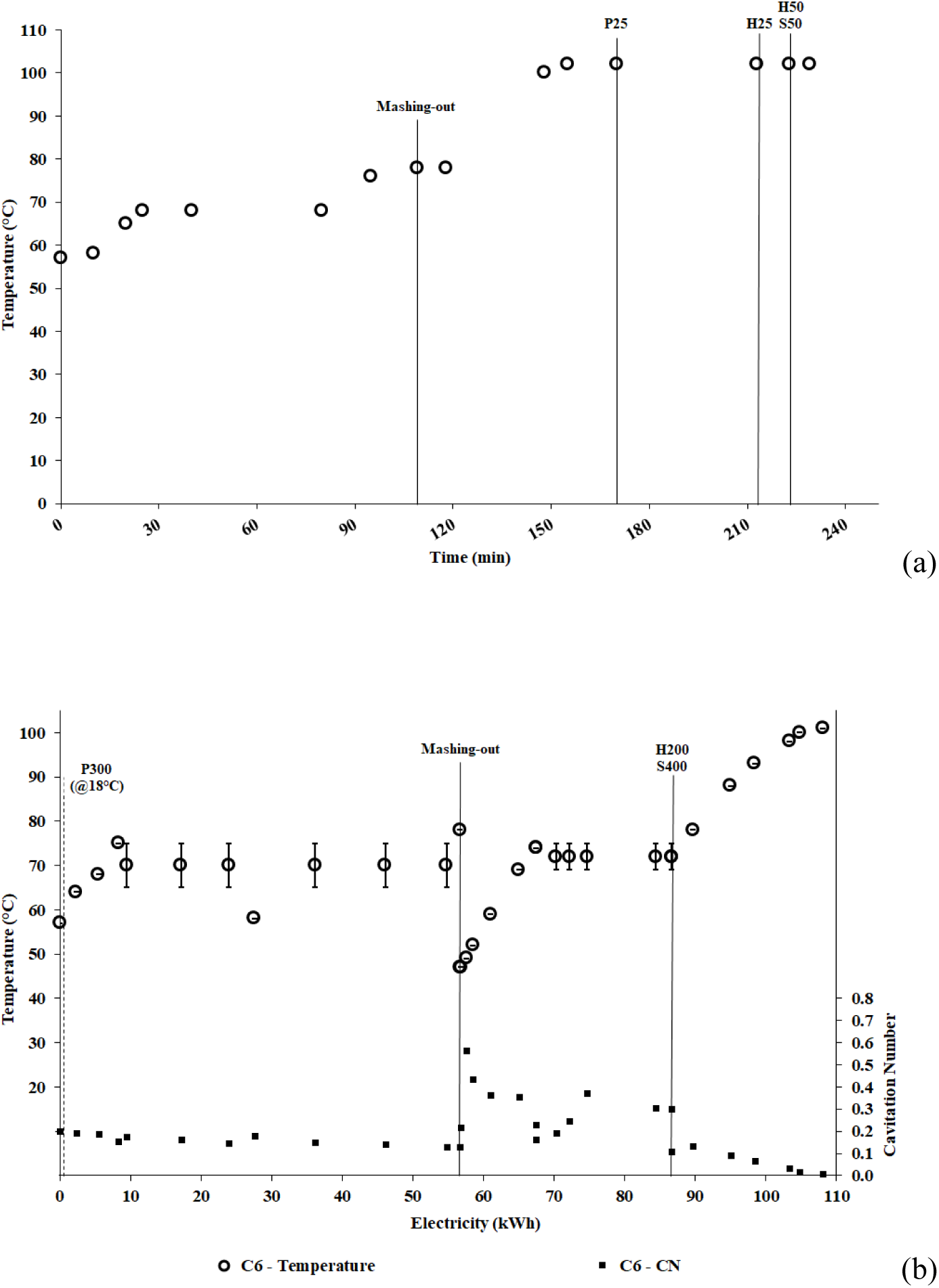
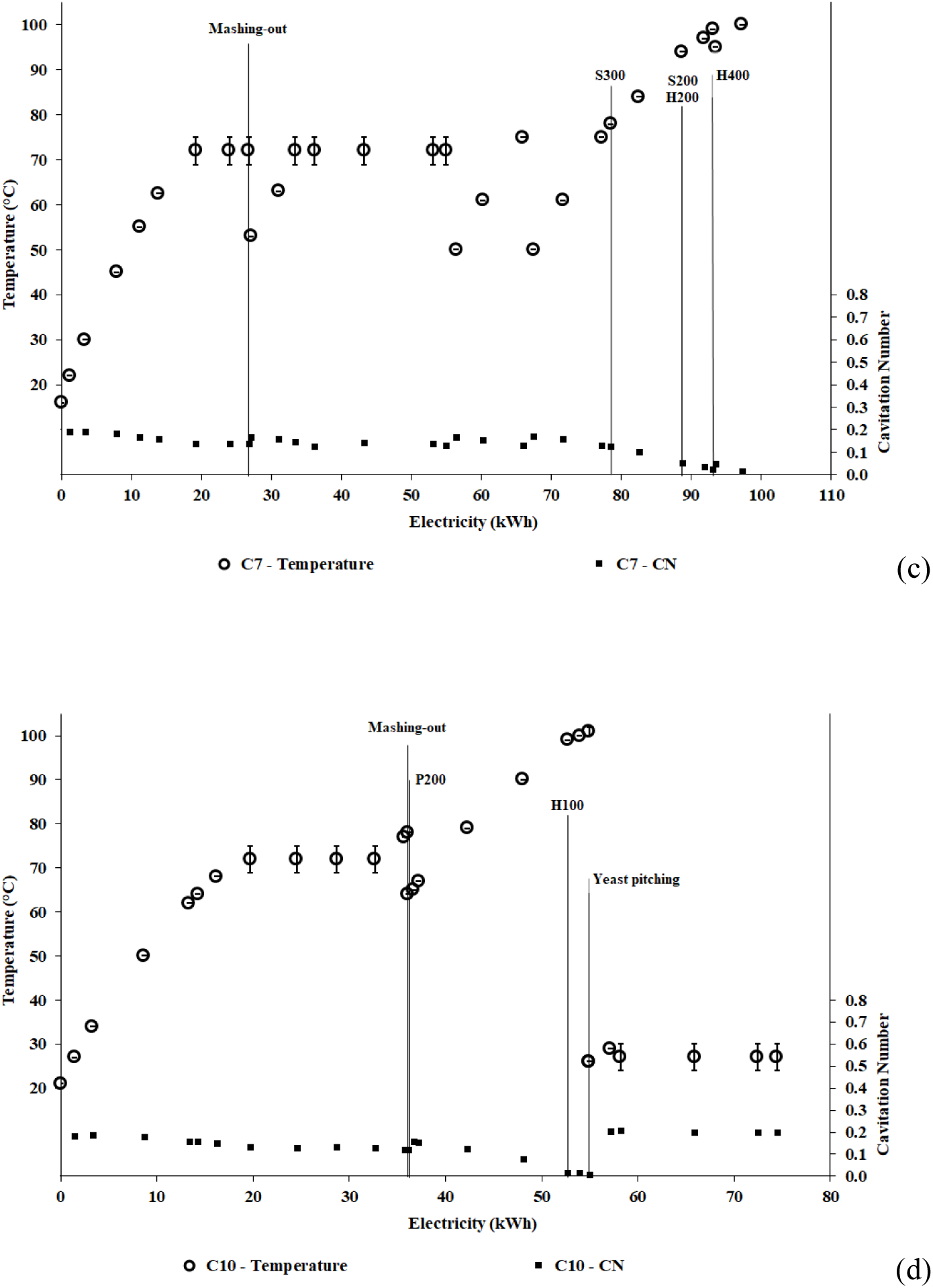
Temperature and basic brewing process stages, as a function of time, for test B1 (a), temperature with respective uncertainties, cavitation number and basic brewing process stages, as a function of consumed electricity, for tests C6 (b), C7 (c), and C10 (d).

Therefore, the different hops batches were inserted in the HC equipment at different moments and phases of the respective processes process times, thus undergoing different cavitation conditions in terms of duration and intensity, as well as immersed in liquids of varying composition. For example, in test C6, a first batch of Perle hops were inserted from the very beginning of water warming (temperature around 18°C), even before malt insertion, while in test C10 all hops underwent cavitation processes together yeasts during about 4 hours.

After fermentation, the analyzed beer samples were bottled by means of an ordinary depression pump, then kept for an average of 4 months at room conditions and, finally, stored during 14 months in a fridge at the temperature of 4 °C. Hydrodynamic cavitation processes could have contributed to increase the shelf-life of the resulting bottled beers, due to the proved complete inactivation of pathogens and undesired microorganisms at moderate temperatures, *i.e.* lower than 60°C (Albanese et al., 2015; Carpenter et al., 2016).

## 3. Results

Table 3 shows that significant differences exist in the amounts of XN, DMX and 6-GN found in the beer samples extracted from tests described in Table 2.

**Table 3.**
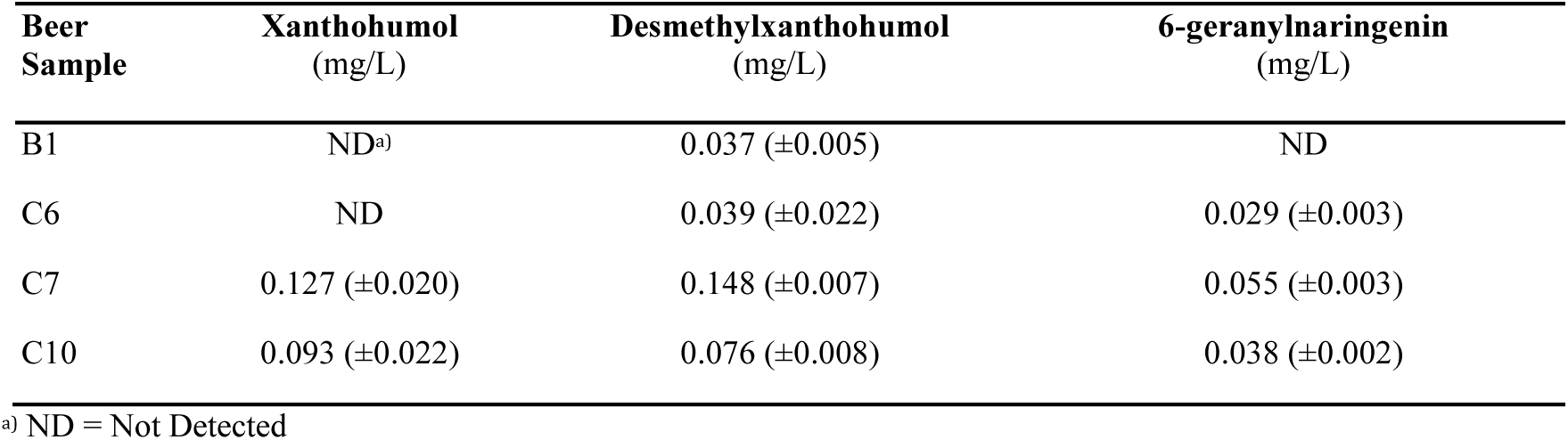
Xanthohumol, desmethylxanthohumol and 6-geranylnaringenin detected in the beer samples shown in Table 1.

The beer B1, obtained via a conventional beer-brewing process, after 18 months did not contain neither XN, nor 6-GN, but only DMX at the level of 37 μg/L. The absence of measurable xanthohumol is surprising, since more than 80% of hops were inserted within 16 min before the end of wort-boiling stage, which should have limited its cyclization to isoxanthohumol. Sample C6, obtained from the test including the longest cavitation process, most of which carried out after malt removal (mashing-out), also contains no xanthohumol. That beer shows a concentration of desmethylxanthohumol comparable to the conventional beer sample B1 (39 *vs.* 37 μg/L), even if with a much larger uncertainty, as well as shows a measurable concentration of 6-geranylnaringenin (29 μg/L). The latter, as recalled from Section 1, is mostly generated in the boiling wort from conversion of the 3′-GCN geranylchalcone found in the hops, therefore cavitation looks like to generate more 6-GN than conventional wort-boiling.

The beer obtained in test C10 does contain xanthohumol (93 μg/L), as well as its concentration of desmethylxanthohumol almost doubles to 76 μg/L in comparison with sample C6, and a significant concentration of 6-geranylnaringenin is also observed (38 μg/L).

A dramatic increase in the concentration of all three prenylflavonoids is observed in the beer sample C7, *i.e.* XN, DMX and 6-GN in the amounts of 127 μg/L, 148 μg/L, and 55 μg/L, respectively.

Fig. 4(a-c) show the results listed in Table 3, compared with published concentrations for Ale-style beers. Concentrations of XN and 6-GN in Ale beers were drawn from the Phenol-Explorer database (Rothwell et al., 2013), while no published values are available for DMX since it is generally absent from final beers (Stevens & Page, 2004).

**Figure 4.**
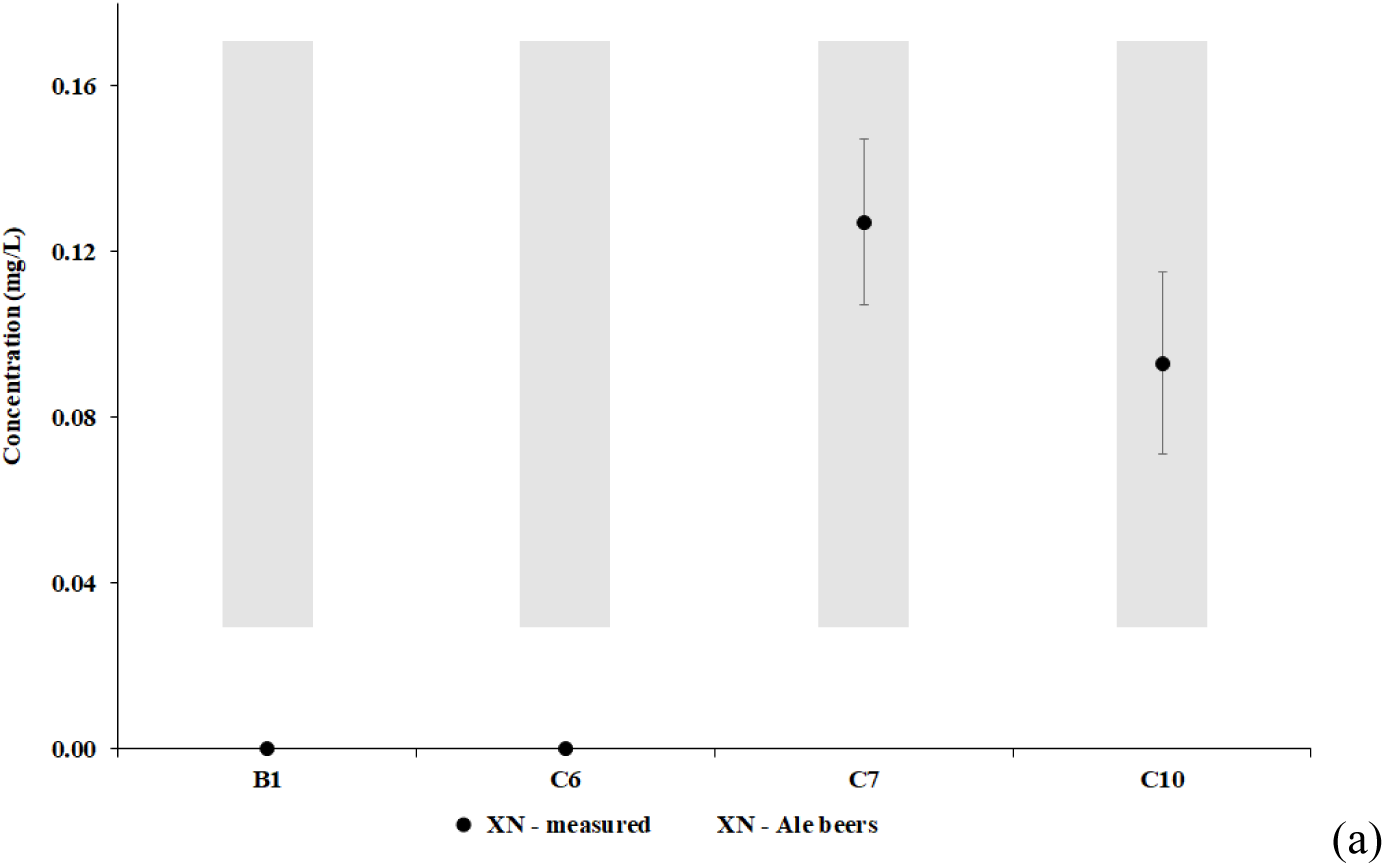
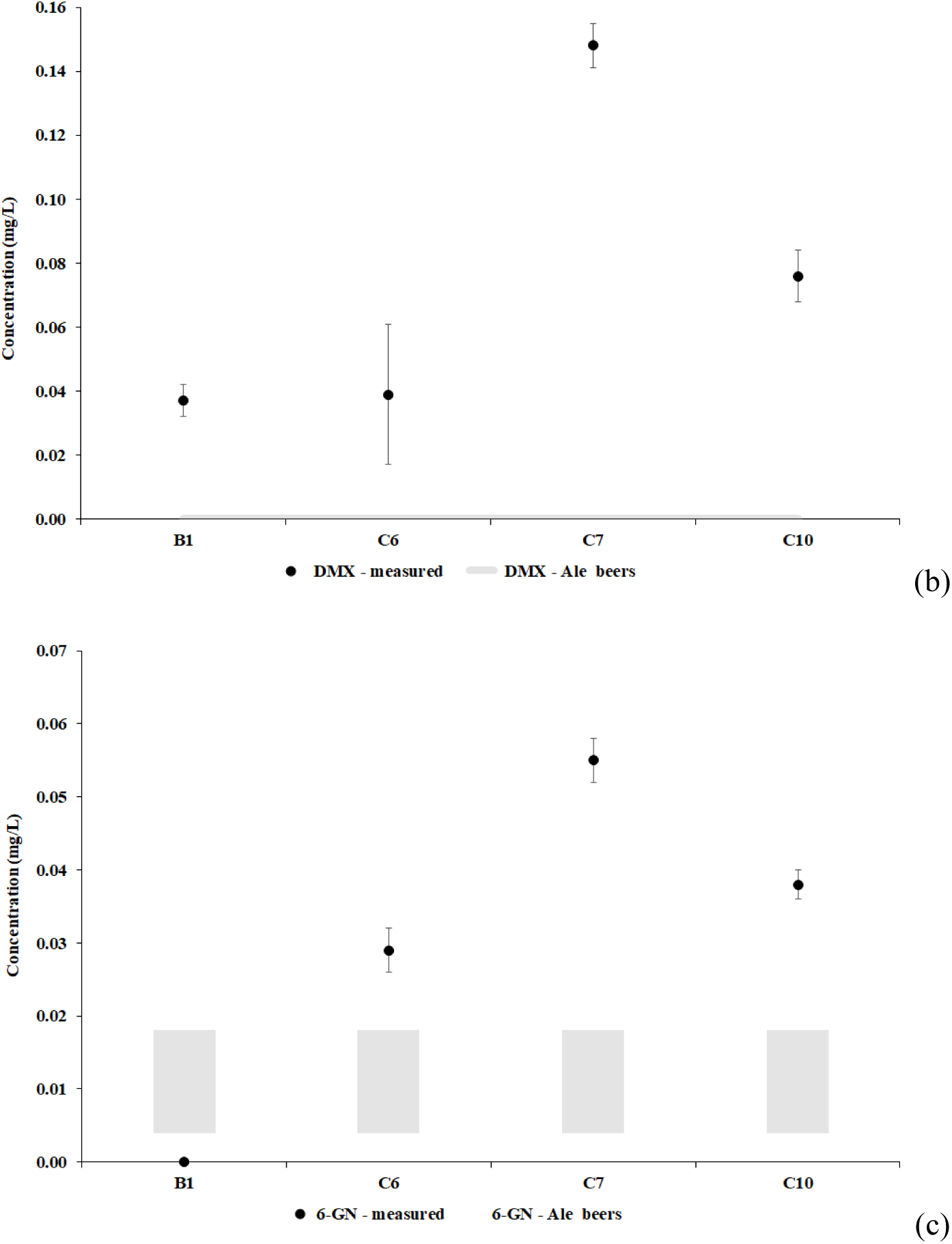
Concentration of xanthohumol (a), desmethylxanthohumol (b), and 6-geranylnaringenin (c) in the analyzed beer samples, compared with the range of values found in Ale-style beers, the latter shown as vertical gray shadows (average values ± standard deviations).

It is observed from Fig. 4(a) that XN concentrations in samples C7 and C10 fall into the range of published values, with the value for the C7 sample in the upper range.

Concentrations of DMX, as shown in Fig. 4(b), are quite surprising, as that molecule is hardly found in final beers due to its thermal isomerization in the brew kettle (Stevens & Page, 2004). Likely, the small but measurable value found for sample B1 derives from the very late insertion of most hops in that brewing process (6 min before the end of the boiling stage), as shown in Fig. 3(a). However, the high concentration found in sample C7 can hardly be explained based on the last insertion of Hers hop (400 g), since it occurred 41 min before the end of the respective HC-assisted brewing process, subject to temperatures in the range 95 to 100°C, as shown in Fig. 3(c). Finally, in test C10, the last hop insertion (Hers, 100 g) was performed 17 min before the end of the warming phase, during such period subject to temperatures between 99 and 101°C. Subsequently, after yeast pitching, hops underwent a long cavitation process (about 4 hours), at low temperatures levels (about 27°C), as shown in Fig. 3(d). Therefore, it appears that low temperature HC processes are, at least partially, unable to convert DMX into a mixture of 6- and 8-prenylnaringenin, the latter a conversion process known to occur in the boiling wort.

Fig. 4(c) shows that the concentrations of 6-GN in all the samples produced from HC-assisted brewing process, *i.e.* samples C6, C7, and C10, exceed the respective concentration range found in Ale beers. The measured level in sample C7 falls in the upper range of the substantially higher values found in Dark beers (Rothwell et al., 2013; Stevens, Taylor, & Deinzer, 1999). Along with the absence of measurable 6-GN in the sample from the conventional test B1, the above observations suggest that, under certain conditions, HC is quite effective in the extraction of 3′-GCN from hops, and/or in its conversion to 6-GN in the processed wort.

## 4. Discussion

The first basic conclusion, so far, is that, under certain conditions, HC-assisted brewing is able to retain the most important of the hops’ prenylflavonoid, *i.e.* xanthohumol, in final beers, at concentrations common to conventionally produced beers, but much greater than the concentration obtained in the performed conventional test (actually, zero), according to measurements following 18 months of beer storage in varying conditions.

As a further important result, it appears that, under the same conditions, HC-assisted brewing is able to retain another hops’ prenylflavonoid, desmethylxanthohumol, significantly better than conventional brewing processes. The common behavior for XN and DMX is hardly surprising, however, based on the common losses and conversion mechanisms mentioned in Section 1, *i.e.* incomplete extraction from hops in the wort, adsorption to insoluble malt proteins, and adsorption to yeast cells during fermentation, besides isomerization in the boiling wort. An hypothesis can be advanced, that HC-assisted brewing helps retaining XN and DMX due to the absence of the boiling stage, the effective extraction from hops – already proved for hops’ α-acids (Albanese et al., 2017a) – as well as, likely, the enhanced solubilization of certain malt proteins, the latter hypothesized in a previous study (Albanese et al., 2017b).

Aiming to provide further inside, Fig. 5(a-c) show the results listed in Table 3, along with the input concentrations of the considered prenylflavonoids in the wort, deriving from the inserted hops as per Table 2 and based on hops’ data listed in Table 1. Vertical scales are very different except for Fig. 5(b).

**Figure 5.**
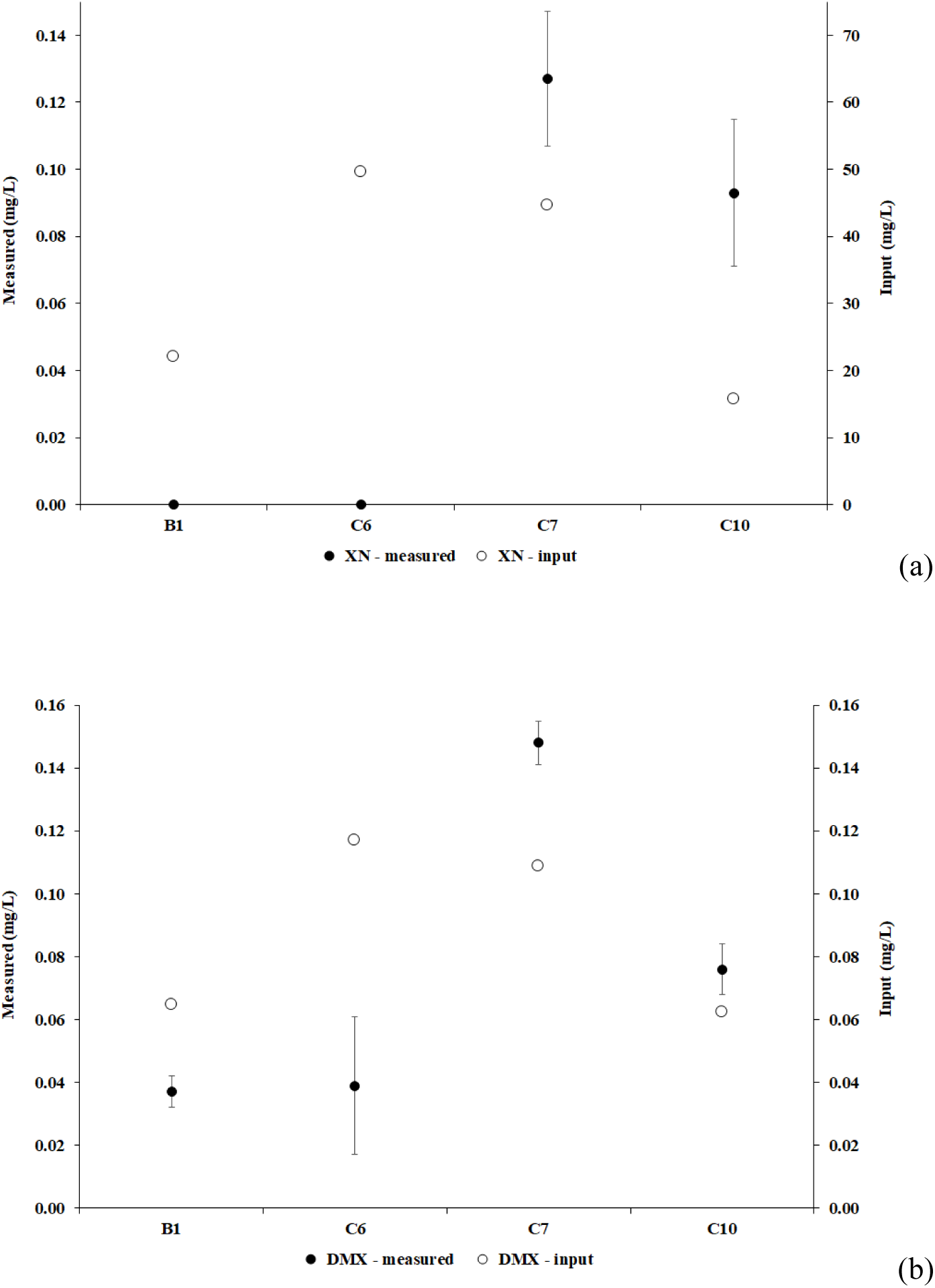
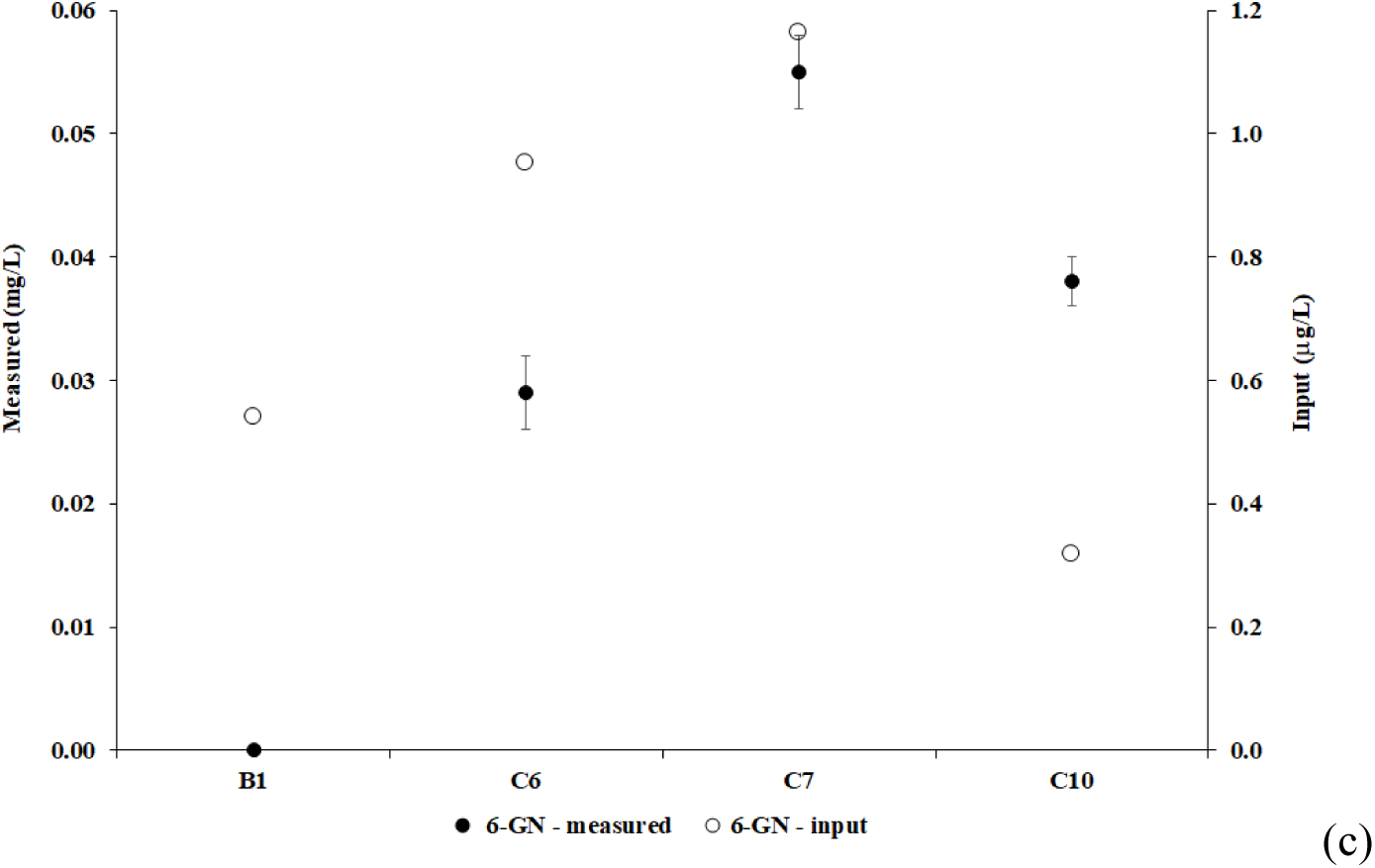
Concentration of xanthohumol (a), desmethylxanthohumol (b), and 6-geranylnaringenin (c) in the analyzed beer samples, along with the input concentrations of the considered prenylflavonoids in the wort.

Fig. 5(a) shows that in beer sample C7, with the greatest absolute XN concentration, the latter is just 0.3% of its input value. Moreover, beer sample C10 retains by far the greatest amount of XN in comparison to the respective input, *i.e.* 0.6%. Comparing with results from the other tests, and following considerations formulated in Section 3 about the respective timing and conditions of hops insertion, preliminary conclusions can be drawn that hydrodynamic cavitation enhances XN extraction from hops and likely reduces its adsorption to insoluble malt proteins, conditioned upon short residence time of hops during cavitation at high temperature levels. In particular, despite the few available data, it looks like hops’ residence times at temperatures above about 80°C, during cavitation, should be less than about 40 min, with better results around 15 to 20 min, while a lower limit on time residence cannot be inferred from the available data. At much lower temperatures, *i.e.* less than 30°C as in test C10 (Fig. 3(d), cavitation does not seem to affect XN concentrations at a significant level, irrespective of its duration. Therefore, HC-assisted brewing can help producing beer higher in XN concentration, conditioned upon inserting hops with higher XN specific content (such as Perle or Saaz, as per Table 1) as late as possible in the hot wort, as well as terminating the process as soon as the desired level of iso-α-acids concentration has been achieved (Albanese et al., 2017a). Moreover, the results of this study suggest that HC-assisted brewing could help the efforts to increase the XN concentration in final beers, such as by means of XN-enriched hops extracts, due to likely partial solubilization of malt proteins.

A more surprising pattern arises from Fig. 5(b). Both beer samples C7 and C10 show DMX concentrations higher than the respective input values, at the levels of 136% and 122%, respectively. While such increase could be merely the result of inherent uncertainties affecting the hops’ DMX concentration values (with likely contribution to uncertainties from hops’ age and breeding conditions), the contrast with sample B1 (57% of input concentration) and sample C6 (33% of input concentration) is striking. The same cavitation conditions inferred for superior retaining of significant concentrations of XN, appear to work much better with DMX, whose cyclization to 6- and 8-prenylnaringenin, described in Section 1, could have been prevented, even if the specific mechanism remains unknown. As well, it is not clear why test C7 produced slightly better results, in relative terms, compared with test C10. Again, the partial solubilization of malt proteins could have played a role in preserving DMX concentrations, the latter likely more effective in the longer test C7. As well, the different main malts used in test C7 (Pilsner) and in test C10 (Pale) could have affected the DMX adsorption.

Finally, Fig. 5(c) shows that 6-GN concentrations in final beers produced by means of HC-assisted brewing processes are much greater than the respective input values, *i.e.* 30 times in sample C6, 47 times in test C7 and as much as 120 times in test C10. This evidence confirms that 6-GN results largely from the conversion of other compounds in the processed wort, namely hops’ 3′-GCN, as explained in Section 1.

Aimed at supplying a raw assessment of the conversion rates from hops’ 3′-GCN to beers’ 6-GN, its average content of 1% of total resin flavonoids for several hops varieties has been assumed, and compared with the average content of 87% for XN (Stevens, Ivancic, Hsu, & Deinzer, 1997). Therefore, 3′-GCN concentration for each hop variety used in the tests has been assumed as 1.5% of the respective XN concentration as per Table 1.

Fig. 6 shows the concentrations of 6-GN listed in Table 3, along with the input concentrations of 3′-GCN in the wort, in different vertical scales.

**Figure 6.**
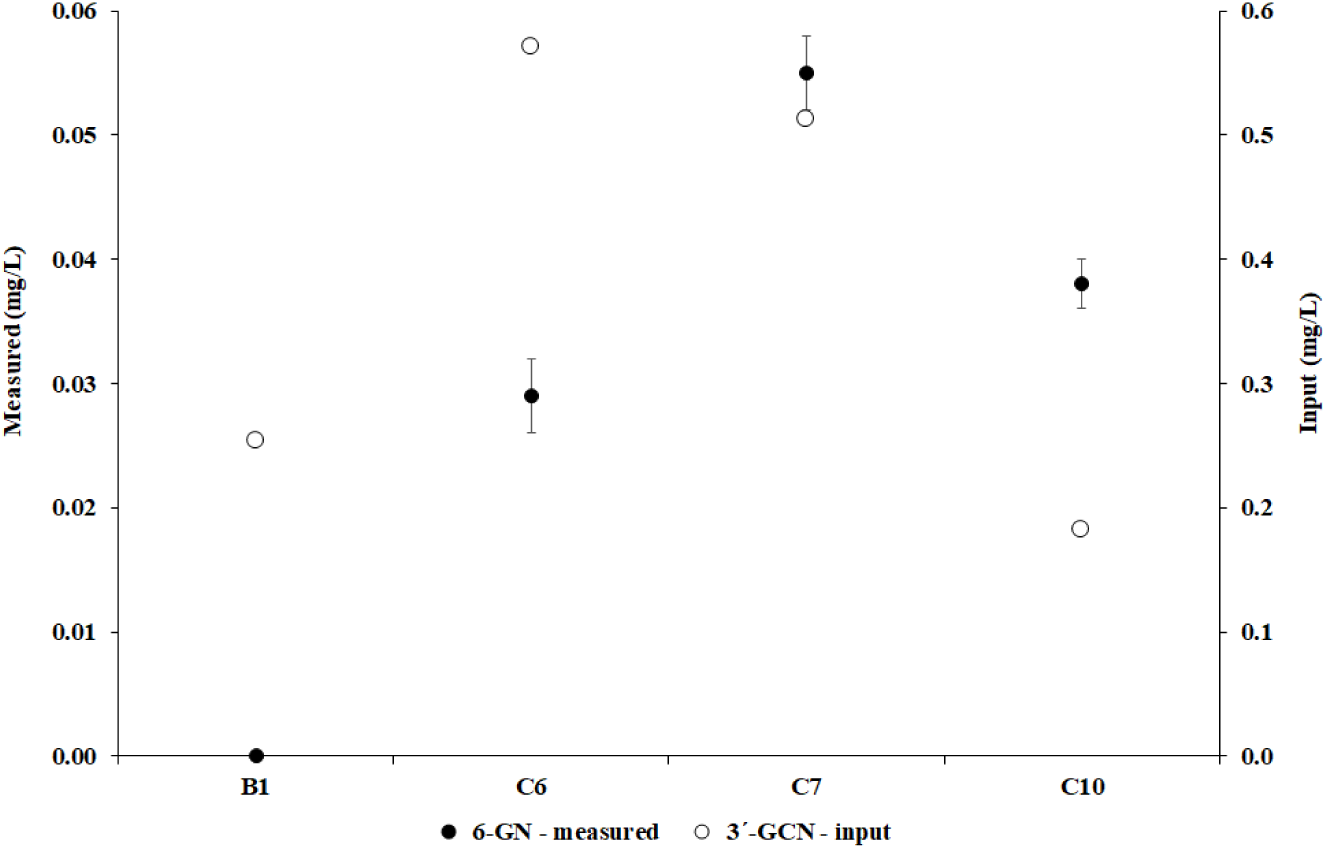
Concentration of 6-geranylnaringenin in the analyzed beer samples, along with the input concentration of its precursor 3′-geranylchalconaringenin in the wort.

Based on Fig. 6, the ratio of 6-GN concentrations in final beers to the input 3′-GCN concentration, change from 0% in the conventional test B1, to 5.1% in C6, 10.7% in C7, up to 20.9% in C10.

The absence of 6-GN in sample B1, the much higher values in samples C6, C7, and C10, in comparison with Ale beers, as shown in Fig. 4(c), as well as the data from Fig. 6 and following discussion, suggest that HC processes are likely allowing a far more effective extraction of the 3′-GCN geranylchalcone from hops, which is the precursor of 6-GN, in comparison to traditional brewing processes.

The relative superiority of test C10, in comparison to the other HC-assisted brewing tests, suggests that the optimal cavitation regimes inferred for retaining XN and DMX can apply to the generation rate of 6-GN in the processed wort (and extraction of 3′-GCN from hops). That is, hops’ residence times at temperatures above about 80°C, during cavitation, should be less than about 40 min, with significantly better results around 15 to 20 min, while a lower limit on time residence cannot be inferred from the available data.

The novelty of the HC-assisted brewing, along with the promising results of the few experiments carried out and discussed in this study, open the way to plenty of further research on the fate of bioactive products from hops to the final beer. The analysis of the transformation and resulting concentrations of XN, DMX and 6-GN compounds under a much broader spectrum of hops (variety and concentration), malts (variety), and HC-assisted brewing conditions (*e.g.*, timing of hops insertion), will represent straightforward continuation of the research presented in this study.

The viability of XN-enrichment of beers, the transformation of many other, minor constituents in hops’ resins (Taniguchi et al., 2014), as well as the respective impact on beer quality, shelf life, and healthy properties, represent some of the open research issues under the new HC-assisted brewing paradigm.

As a final remark, the findings of this study agree with recent literature, showing the viability of hydrodynamic cavitation as an effective, efficient and scalable technology to boost the extraction of food bioactive compounds (Roohinejad et al., 2016), as well as to release bound phenolics and increase the antioxidant activity of processed food (Lohani, Muthukumarappan, & Meletharayil, 2016).

## 5. Conclusions

Based on the results of preliminary tests, HC-assisted brewing appears able to retain, or generate, higher concentration of certain important hops’ bioactive compounds in the final beer, such as the prenylflavonoids xanthohumol, desmethylxanthohumol, and 6-geranylnaringenin. Besides allowing cheaper, more efficient, safe and reliable, and more environmentally friendly processes (Albanese et al., 2017a), as well as affording substantial gluten reduction without additives or complex technological processes (Albanese et al., 2017b), hydrodynamic cavitation was proven a viable way to produce healthier beers, exploiting hops’ unique natural products. An with the gluten reduction issue, optimal cavitational and process conditions, aimed at retaining and generating the hops’ bioactive compounds under consideration, were preliminarily identified, even if much research remains to be done to explore a wider range of beer recipes, and fully exploit the promising capabilities of HC-assisted brewing, including the use of different cavitation reactors.

## Acknowledgements

L.A. and F.M. were partially funded by Tuscany regional Government under the project T.I.L.A. (Innovative Technology for Liquid Foods, Grant N°. 0001276 signed on April 30, 2014). Part of this research was carried out under a cooperation between CNR-IBIMET and the company Bysea S.r.l., with joint patent submitted on August 9, 2016, international application No. PCT/IT/2016/000194 “A method and relative apparatus for the production of beer”, pending.

## Declaration of interest

L.A. and F.M. were appointed as Inventors in the patent submitted on August 9, 2016, international application No. PCT/IT/2016/000194 “A method and relative apparatus for the production of beer”, pending.

